# A 7-channel high-*T*_c_ SQUID-based on-scalp MEG system

**DOI:** 10.1101/534107

**Authors:** Christoph Pfeiffer, Silvia Ruffieux, Lars Jönsson, Maxim L. Chukharkin, Alexei Kalaboukhov, Minshu Xie, Dag Winkler, Justin F. Schneiderman

## Abstract

Due to their higher operating temperature, high-*T*_c_ superconducting quantum interference devices (SQUIDs) require less thermal insulation than the low-*T*_c_ sensors that are utilized in commercial magnetoencephalography (MEG) systems. As a result, they can be placed closer to the head, where neuromagnetic fields are higher and more focal, potentially leading to higher spatial resolution. The first such on-scalp MEG measurements using high-*T*_c_ SQUIDs have shown the potential of the technology. In order to be useful for neuroscience and clinical applications, however, multi-channel systems are required. Herein, we present a 7-channel on-scalp MEG system based on high-*T*_c_ SQUIDs. The YBCO SQUID magnetometers are arranged in a dense, head-aligned hexagonal array inside a single, liquid nitrogen-cooled cryostat. The spacing between the magnetometers and the head is adjustable down to 1 mm. The sensors are side-mounted on the cryostat that is mounted on an articulated armature for recordings on arbitrary head locations of a seated subject. We demonstrate white noise levels of 50-130 fT/Hz^1/2^ at 10 Hz, sensor-to-sensor crosstalk values of *<*0.6%, and single-fill operation times of 16 hours. We validate the system with MEG recordings of visual alpha modulation and auditory evoked fields. The system is thus useful for densely and sensitively sampling neuromagnetic fields over any *∼* 10 cm^2^ patch of the scalp surface over the course of a day.

## 1 Introduction

Magnetoencephalography (MEG) is a functional neuroimaging method based on measuring magnetic fields originating from neuronal currents in the brain. Due to its high temporal and spatial resolution, MEG is a useful tool in the effort to better understand the human brain - and as a result learn how to treat it in disease. Because of its unsurpassed combination of spatial and temporal resolutions, MEG is used extensively for studies of sensory processing, language, plasticity, memory encoding, connectivity, development, etc. [1, 2]. In clinical settings, MEG is also increasingly used to localize epileptic foci and for pre-surgical planning via mapping of eloquent regions of the brain [3–5].

Commercially available, state-of-the-art MEG systems employ hundreds of low noise, low-*T*_c_ SQUID sensors. A major limitation of these systems stems from the fact that such low-*T*_c_ SQUIDs are cooled with liquid helium (T ~4.2 K). This extreme cryogenic temperature necessitates more elaborate cryogenics, resulting in a large standoff distance (~20 mm) between the sensors and room-temperature—and consequently to the head of the subject. This standoff distance limits the measureable neuromagnetic signal magnitudes, as magnetic fields decrease with distance from their source.

Alternative sensor technologies that operate at higher temperatures can reduce this sensor-to-source distance. High-*T*_c_ SQUIDs, for example, can operate at the boiling temperature of liquid nitrogen (T ~77 K); the distance for such sensors can thus be reduced to less than 1 millimeter (consisting of vacuum space and a sub-mm thin vacuum-supporting window) [6]. One limitation, however, is that the noise levels of high-*T*_c_ SQUIDs are usually higher than that of their low-*T*_c_ counterparts. This can be partially mitigated by the signal gain and different spatial sampling that stem from coming closer to the head. As such, it is possible to obtain more information from the brain with high-*T*_c_ vs low-*T*_c_ SQUID technology, despite the inferior sensor noise levels of the former [7, 8].

MEG recordings have been demonstrated with single high-*T*_c_ SQUIDs in 1996 [9]; measurements with greatly reduced standoff distance—i.e., “on-scalp” MEG as it is now called—have followed since 2012 [10–12]. Such recordings have verified the expected increase in signal strength when sampling closer to the head. As an important next step, a single high-*T*_c_ SQUID has been used to localize evoked activity with serial recordings of repeated stimulation sessions while the sensor was positioned at different locations on the head [13]. Localizing in such a way is, however, time consuming and limited to evoked activity. The sources of relevant neural responses are furthermore not always known beforehand and may include activity involving several (connected) brain regions. Finally, modern noise reduction techniques utilized in MEG, e.g., Signal Space Separation (SSS) [14], require multi-channel systems (preferably full-head). Simultaneously sampling the magnetic field at many locations on the head is therefore crucial for many modern MEG applications, necessitating the development of multi-channel systems for on-scalp MEG. As a step towards full-head coverage, we have developed a 7-channel high-*T*_c_ SQUID-based on-scalp MEG system.

## 2 Methods

### 2.1 Cryostat

A schematic of the system is presented in Fig. 1. At the center of the cryostat is a 0.9-liter liquid nitrogen (LN_2_) tank made from epoxy-reinforced glass fiber. The thermal insulation for the main body of this tank consists of multiple layers of superinsulation foil within a vacuum space. A charcoal trap attached to the nitrogen tank provides additional cryopumping for the vacuum insulation when the system is cold. A jacket of the same epoxy-reinforced glass fiber material forms the outer shell of the cryostat, providing an enclosure for the vacuum as well as structural support.

**Figure 1:**
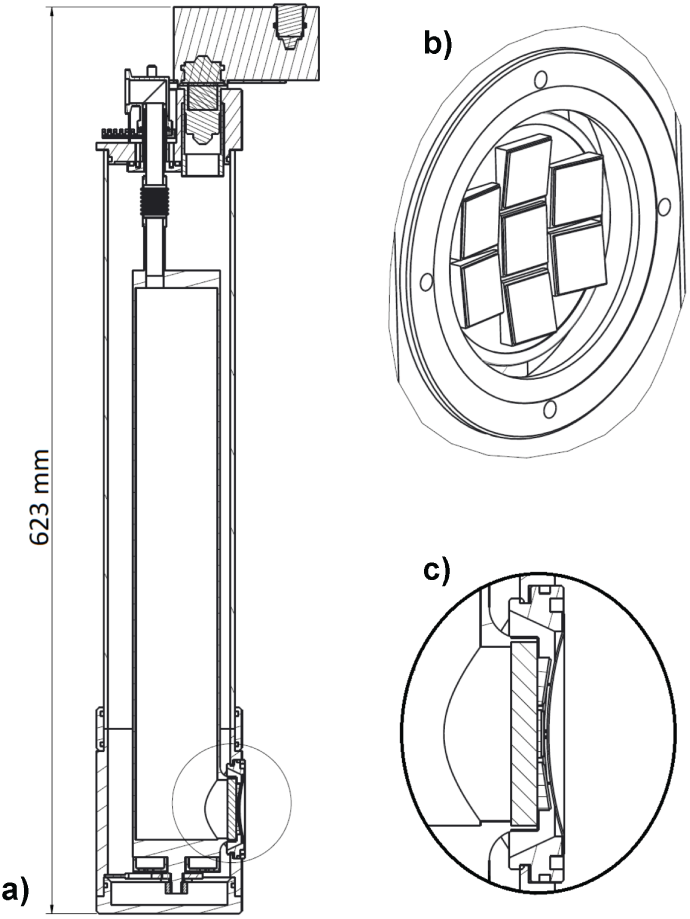
Schematic of the cryostat. a) cross-section of the cryostat. An electronics breakout box is mounted to the top. The LN_2_ transfer and pump port is connected to the top of the LN_2_ vessel via a bellows. b) the sapphire support and wedges holding the SQUID sensors. c) zoom of the support structure for the SQUIDs and window in cross-section

The SQUIDs are mounted on a 44 mm diameter sapphire disk that seals a hole on the side of the LN_2_ tank; this support provides high thermal contact between the sensors and the LN_2_. The side-mounting allows for a wide range of cryostat orientations suitable for neuromagnetic recordings on supine and seated subject positions. Seven 11 mm × 11 mm sapphire wedges that hold the SQUIDs are glued to the sapphire disk in a hexagonal array with one in the center. The sensors are tightly packed (2 mm edge-to-edge) for high spatial sampling of a small patch of the head surface (~ 10 cm^2^). Fig. 1-b shows a drawing of the SQUID array. The outer wedges are titled towards the center, aligning the SQUIDs to the surface of a sphere with a radius of 8 cm, which roughly corresponds to the average curvature of an adult head. A 0.4 mm thick, concave, room-temperature, and vacuum-supporting window is set into the vacuum jacket next to the sensors (Fig. 1-c). Due to the window’s curvature and a screw connection on the window frame that allows adjustment of the distance between window and sensors, a minimal (as low as 1 mm) distance between all of the sensors and the subjects head can be achieved.

### 2.2 High-*T*_c_ SQUID magnetometers

The cryostat houses seven single layer YBa_2_Cu_3_O_7*−x*_ (YBCO) SQUID magnetometers, each of which has a directly coupled pickup loop. The current design is similar to the one our group has used for magnetometers in single channel cryostats [10, 11], but has been adapted to allow for dense sensor packing with on-chip feedback [15]. Each hairpin SQUID includes a pair of grain boundary Josephson junctions and is made on a 10 mm × 10 mm SrTiO_3_ (STO) bicrystal substrate with a misorientation angle of 22.6^*◦*^ (Shinkosha, Japan). The pickup loop has outer dimensions of 8.6 mm × 9.2 mm and a linewidth of either 1 or 3 mm. While the narrow linewidth magnetometers have shown low flux trapping and crosstalk, wider (3 mm) pickup loop lindwidths theoretically have a superior effective area to inductance ratio. As this should result in lower magnetic field noise values for otherwise identical SQUIDs, we made 3 wide pickup loop magnetometers for comparison.

Each sensor is fabricated by sputtering a CeO_2_ buffer layer, which is followed by in-situ deposition of an epitaxial YBCO 150-225 nm thin film with pulsed laser deposition (PLD). The high-*T*_c_ superconducting film is then patterned with a standard photolithography process using a laser writer and argon ion etching. We manually round one edge of the substrate to make contact pads that can be accessed from the side. These side contacts reduce the standoff distance because the cryostat wires no longer protrude into the space between the top surface of the sensor substrates and the vacuum-supporting window. Gold contact pads are made by magnetron sputtering and a lift-off process.

The SQUIDs are operated in a flux-locked loop (FFL) in order to linearize the periodic output signal and increase the dynamic range [16, 17]. We apply the feedback flux by direct injection of current into the SQUID loop. This approach is favorable for densely-packed SQUIDs in an on-scalp MEG system as it eliminates the need for an additional feedback coil i.e., it is a simple on-chip method that compromises neither the standoff distance nor the thermal connection of the sensors to the LN_2_. It furthermore leads to low sensor-to-senor feedback crosstalk (less than 0.5% between adjacent magnetometers) [15]. Three commercial 3-channel direct readout electronics with AC-bias reversal (SEL-1 from Magnicon) are used to control the SQUIDs [17]. The AC-bias reversal mode reduces low frequency 1/f-like noise due to critical current fluctuations, but we find that it couples noise into neighboring sensors. Such bias reversal crosstalk could, however, be mitigated by modifying the electronics such that all channels share a single bias reversal clock. When using synchronized clocks, bias reversal crosstalk induces a constant flux bias for each SQUID, which can be manually compensated. This flux bias is routinely adjusted when operating a single SQUID; for multi-channel operation it is, however, crucial to set the bias currents of all SQUIDs before adjusting the flux biases.

## 3 Results

### 3.1 System

Photographs of the complete system with and without the window are presented in Fig. 2. The breakout box connected to the top end of the cryostat contains shunt resistors and diodes (for protecting all of the channels from electrostatic discharge and other potentially damaging current spikes) as well as connectors for the SQUID electronics boxes.

**Figure 2:**
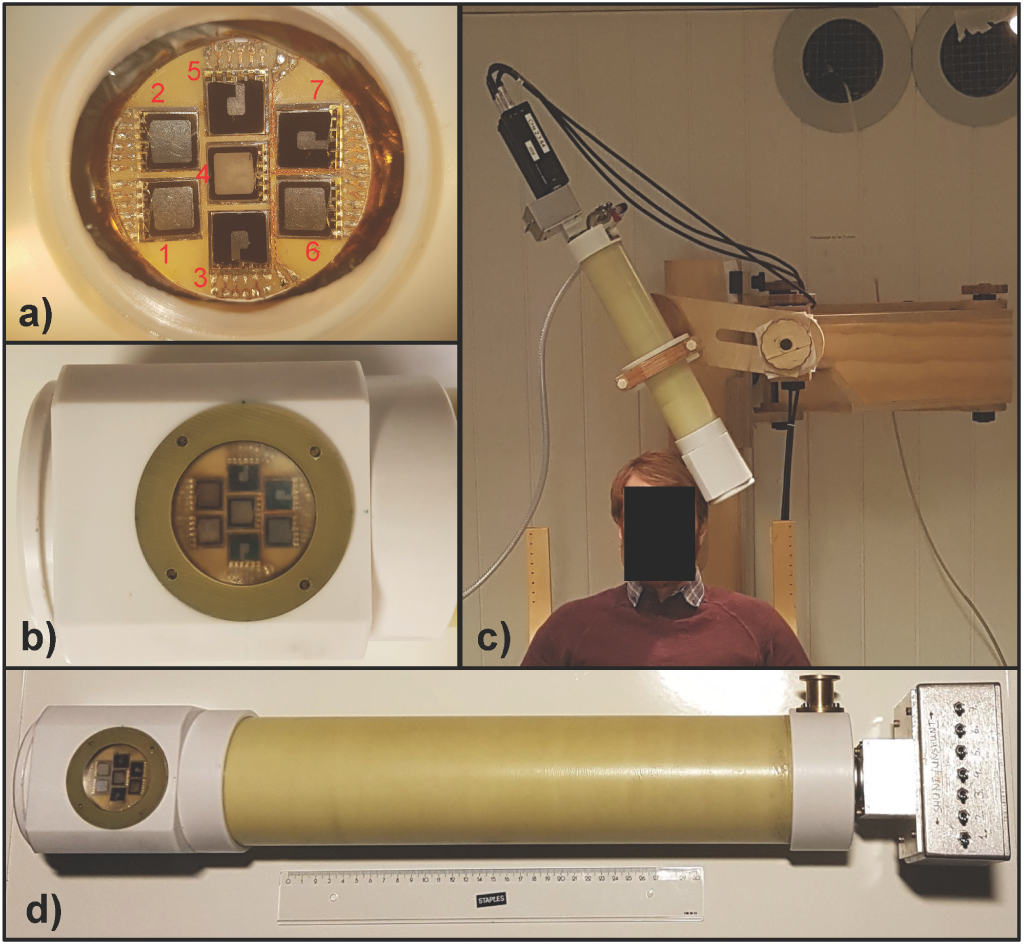
Photographs of the cryostat. a) The sensors mounted on the sapphire wedges and disk (without the vacuum-supporting window). A PCB surrounds the wedges and provides electrical connections to each sensor via low-profile bonds. Channel numbering is shown in red. b) A closeup of the cryostat tail with the side-mounted sensors sitting behind the curved, vacuum-supporting, and height-adjustable window. c) Setup for MEG recordings. The cryostat is supported against the head of a seated subject with a wooden articulated armature. The SQUID electronics and breakout box can be seen connected at the top end of the cryostat. d) Closeup of the complete cryostat with breakout-box.

The sapphire window reaches a base temperature of 80.1 K without pumping on the LN_2_. By reducing the boiling temperature of the LN_2_ via pumping the nitrogen vessel with a pump pressure of ~150 mBar, the temperature on the sapphire support drops to 70.1 K. With a single filling, the cryostat’s hold time (T *<*80 K) amounts to approximately 19 hours without and 16 h with pumping - more than sufficient for a typical day of MEG recordings.

The noise spectra of all seven SQUIDs operating in AC bias reversal mode (40 kHz bias reversal frequency) are presented in Fig. 3. Flat white-noise levels persist down to 6-10 Hz (with a few peaks at 50 Hz and harmonics). With the exception of channel 3, all SQUIDs reach ≤100 fT/Hz^1/2^, with the best channels reaching 50 fT/Hz^1/2^. More details on the magnetometer operation parameters can be found in the appendix A.

**Figure 3:**
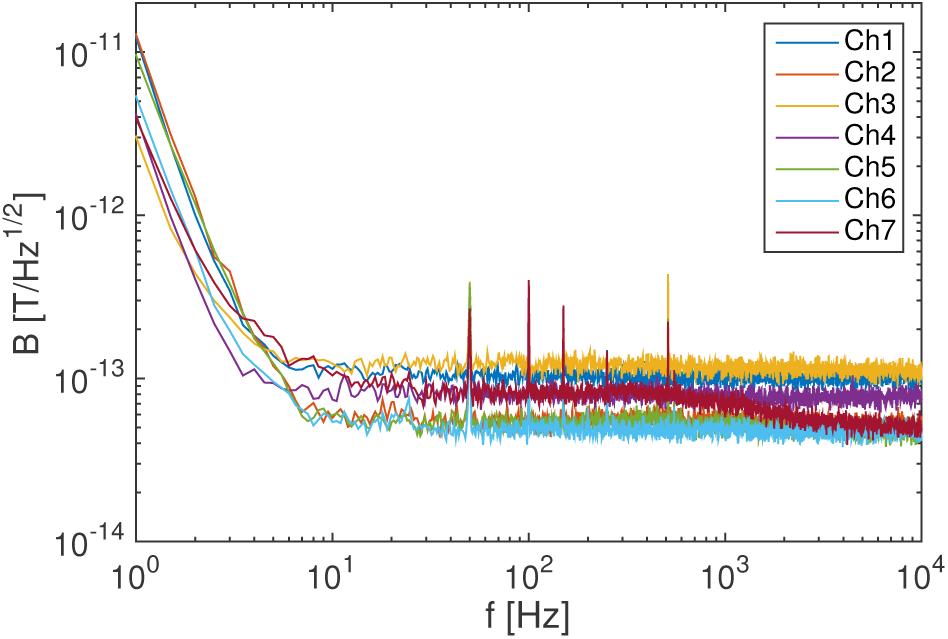
Magnetometer noise spectra recorded in the 7-channel system inside a magnetically shielded room with 2 layers of mu-metal.

Feedback flux crosstalk between all pairs of magnetometers was characterized by applying a sinusoidal current corresponding to 1 flux quantum to the feedback coil of an exciting magnetometer in open loop configuration and measuring the amount of flux coupled into all other sensors. Three magnetometer pairs show elevated crosstalk: the central sensor (channel 4) induces 0.59% in channel 6 and 0.40% in channel 3, while channel 7 induces 0.46% into channel 6. As these values are far larger than those of magnetometer pairs with equivalent relative positioning, we conclude that the main part of this crosstalk is not due to direct coupling of the on-chip feedback to the pickup loops. The increased crosstalk is more likely caused by the twisted wire pair (which is used for supplying the feedback current to the central sensor) that passes between channels 3 and 6. The summed feedback flux crosstalk in the central sensor (which is expected to accumulate the most direct coupling crosstalk due to its close proximity to all other sensors) is 0.22%; for the remaining channels (excluding 3 and 6) it is below 0.2%. The full crosstalk matrix can be found in the appendix B.

### 3.2 MEG recordings

All recordings were performed inside a magnetically shielded room with 2 layers of high permeability nickel-iron alloy (mu-metal) and one layer of copper-coated aluminum (Vacuumschmeltze GmbH). A wooden fixture with a movable articulated armature held the cryostat in place on the seated subject’s scalp during the sessions (see Fig. 2-c). The SQUID signals were digitized with a PC-controlled 16-bit DAQ (NI-USB6259, National Instruments); a second PC was used for communication with the SQUID control electronics unit. The recorded data was subsequently analyzed in Matlab (Matlab R2015a, Mathworks, Natick, MA, USA) using the FieldTrip toolbox [18]. A second DAQ (NI-USB6258, National Instruments) connected to the electronics control computer generated stimulation and trigger signals. The experiment was conducted in compliance with national legislation and the code of ethical principles (Declaration of Helsinki).

#### 3.2.1 Induced alpha activity

We measured modulated alpha rhythm activity (8-12 Hz, [2]) in a single subject. The subject was instructed to fixate gaze upon the center of a wooded nature scene and alternate (upon an auditory cue) between opened and closed eyes (30 seconds each) for a total of 30 minutes. The sensors were aligned to receive signals from the visual cortex by placing the cryostat window against the occipital part of the subject’s head (centered between O1 and O2 in the 10-20 reference system [19]). The sensor signals were bandpass-filtered (1 to 1000 Hz) and amplified (gain of 200) with a custom 7-channel amplifier system developed and built for the cryostat. The data was afterwards cut into epochs (20 seconds before to 20 seconds after opening of the eyes). Time-frequency spectra of the epochs were then calculated and averaged. We present the resulting time-frequency spectra for all 7 channels averaged over 30 trials in Fig. 4. As expected, a suppression of alpha activity was observed after the subject opened the eyes. The signal-to-noise ratio was furthermore high enough for the alpha modulation to be observed at the single-trial level (see Fig. 5).

**Figure 4:**
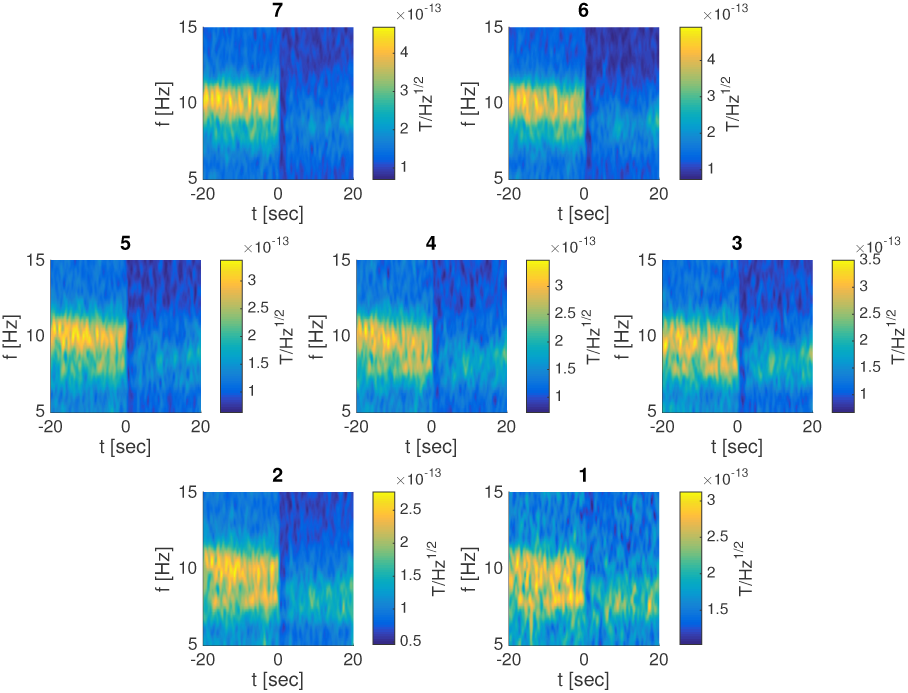
Averaged time-frequency spectra of all 7 channels. Suppression of alpha activity (8-12 Hz) can be seen upon opening of the eyes (at t=0).

**Figure 5:**
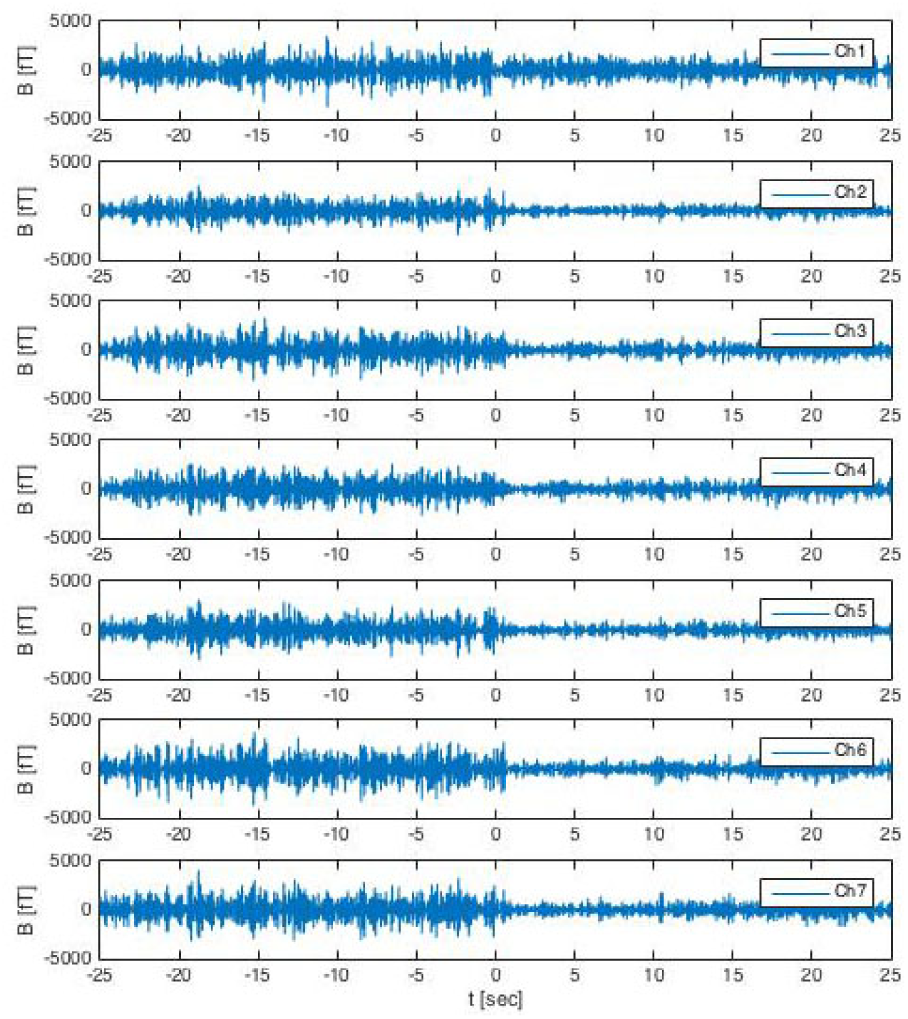
Single trial of alpha suppression (bandpass filtered between 5 and 15 Hz). A decrease in amplitude can be observed after the subject opens the eyes (at t=0).

#### 3.2.2 Evoked auditory activity

We also measured evoked auditory activity in a single subject. The subject was presented with a tone (1000 Hz, lasting 400 ms) every second (with ±10 ms added jitter). The cryostat was placed with the sensors at the left temporal part of the subject’s head (centered on T3 in the 10-20 reference system). To avoid habituation, 1 out of each set of 5 stimuli were randomly replaced by oddball tones (1200 Hz). The tones were generated outside the magnetically shielded room and transferred to the right ear using a plastic ear-tube. As with the alpha recording, the sensor signals were bandpass-filtered (1 to 1000 Hz) and amplified (gain of 200) with our 7-channel amplifier system. An additional Chebyshev bandpass-filter (1 to 200 Hz) was applied to the recorded data before epoching (−100 to 700 ms), and averaging (with removal of a baseline period from −100 to 0 ms) the trials. Fig. 6 includes the resulting time-locked averages of 1000 trials for all 7 channels. A strong N100m peak (at t ~100 ms after stimulus onset [20]) is clearly visible in all channels.

**Figure 6:**
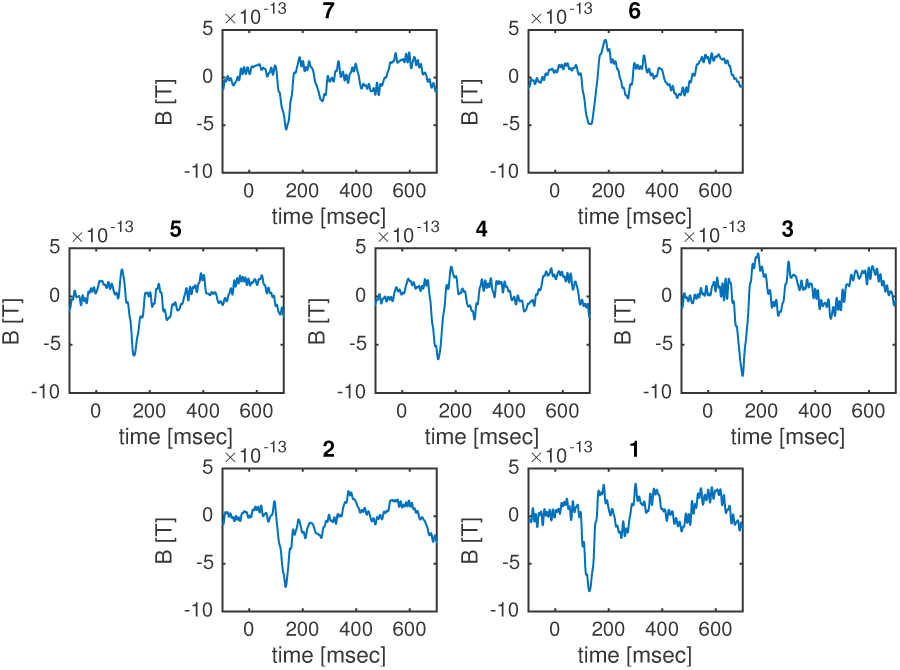
Time-locked averages of auditory evoked fields with the 7-channel on-scalp MEG system. All seven channels show typical auditory N100m response.

## 4 Discussion

While on-scalp MEG with multiple single-channel units has been demonstrated with both high-*T*_c_ SQUIDs [10] and optically pumped magnetometers (OPMs) [21, 22], the only previous report of a multi-channel system is with the latter technology [23]. Due to inherent advantages and disadvantages of high-*T*_c_ SQUIDs compared to OPMs as well as specific design goals described below, our system differs in several key aspects from such on-scalp MEG systems.

Based on experience with single channel recordings, as well as theoretical considerations, the 7-channel on-scalp MEG system described here was designed with two main goals in mind: small sensor-to-head standoff distance and high spatial sampling. The former in order to measure as close to “on-scalp” as possible and the latter because coming closer to the neuromagnetic sources results in higher spatial frequencies contained in the neuromagnetic fields. Higher spatial frequencies potentially lead to better spatial resolution, but also require denser spatial sampling. With 12.0 mm vertical and 13.4 mm diagonal center-to-center distances between adjacent sensors, the spatial sampling of the system presented here is higher compared to that of previously reported multi-channel on-scalp MEG systems (18 mm within 4-channel OPM in [23]; not specified in [21,22], but necessarily greater than the 13 mm x 19 mm sensor footprint [24]). Furthermore the sensor-to-head distance, ranging between 1 and 3 mm, is significantly smaller than in previously reported systems (12 mm in [23] and 6.5 mm in [21, 22]).

High-*T*_c_ SQUIDs have inherently higher dynamic range and bandwidth than OPMs. The 7-channel on-scalp MEG system can therefore be operated inside a standard MSR (we operated it inside a MSR with 2 layers of mu-metal) without any additional shielding or field compensation systems [21, 22]. The bandwidth of the system is furthermore limited only by the SQUID electronics (in our case, where we used a 40 kHz AC-bias, this would be ~20 kHz).

We intend to use the system in further MEG experiments, where we will compare it to conventional, full-head MEG systems as well as investigate new research questions. The combination of measuring closer to the scalp and sampling at higher spatial density is expected to lead to an increase in resolution for on-scalp MEG [25, 26]. This could allow us to observe previously unseen activity. Due to the limited coverage, however, great care has to be put into the design of experiments. In order to capture all relevant field components of a given neural activity, smaller systems, like the one presented here, may have to rely on combining multiple, consecutive recordings requiring accurate co-registration.

## 5 Conclusion

We have developed a multi-channel on-scalp MEG system employing seven high-*T*_c_ SQUID magnetometers in a common cryostat with a single-fill operation time of 16 hours. The sensors are arranged in a dense (2 mm edge-to-edge), head-aligned array with an adjustable separation to room temperature of down to 1 mm. The magnetometers reach noise-levels of 50-130 fT/Hz^1*/*2^ at 10 Hz and show sensor-to-sensor crosstalk below 0.6%.

The system has been validated with MEG measurements of alpha activity and auditory evoked fields that show high potential for future neuroscience experiments.

### A High-*T*_c_ SQUID operation parameters

The high-*T*_c_ SQUID parameters of a typical measurement session are summarized in Table 1. We cool the SQUIDs such that the highest bias current *I_b_* remains below the limit of the electronics (i.e. 250 *µ*A). The bias currents are tuned with the aim of maximizing the voltage modulation depth ∆*V* of the SQUIDs.

**Table 1:** High-*T*_c_ SQUID operation parameters

|  | Ch1 | Ch2 | Ch3 | Ch4 | Ch5 | Ch6 | Ch7 |
| --- | --- | --- | --- | --- | --- | --- | --- |
| $I_b$ [μA] | 3 | 75 | 236 | 159 | 125 | 49 | 67 |
| $\Delta V$ [μV<sub>pp</sub>] | 9 | 48 | 33 | 46 | 38 | 39 | 43 |
| $I_f$ [μA] | 31.6 | 31.8 | 43.7 | 29.4 | 48.9 | 39.0 | 40.3 |
| $L_c$ [pH] | 65.5 | 65.1 | 47.3 | 70.4 | 42.3 | 53.1 | 51.4 |
| $A_{eff}$ [mm<sup>2</sup>] | 0.270 | 0.270 | 0.243 | 0.303 | 0.217 | 0.211 | 0.257 |
| $A_{eff}^{-1}$ [nT/$\Phi_0$] | 7.67 | 7.67 | 8.53 | 6.83 | 9.56 | 9.82 | 8.06 |
| $A_p/L_p$ [mm/nH] | 4.12 | 4.15 | 5.13 | 4.31 | 5.12 | 3.97 | 5.00 |

The feedback current *I_f_* necessary to couple one flux quantum into the SQUID is used for crosstalk measurements. We furthermore estimate the coupling inductance *L_c_* of the pickup loop to the SQUID loop using *L_c_* = Φ_0_*/I_f_*.

We measure the magnetometer effective area *A_eff_* by applying a calibrated magnetic field to the sensor with a Helmholtz coil. The effective area and its inverse, the flux-to-field transfer coefficient, can be used to convert the measured flux to magnetic field units. With the help of the magnetometer effective area *A_eff_* and the coupling inductance *L_c_*, we can estimate the pickup loop effective area to inductance ratio *A_p_/L_p_*:

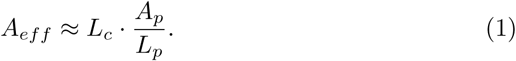

We see that this ratio is on average 23% bigger for the magnetometers whose pickup loops were wide (3 mm: channels 3, 5 and 7) than for the narrow (1 mm width pickup loop) ones.

### B Feedback flux crosstalk

Table 2 shows the flux crosstalk induced from an exciting magnetometer (rows) to a sensing magnetometer (columns). Crosstalk values above 0.2% are marked bold. We furthermore provide sums of crosstalk values for each sensor to estimate the total flux each sensor receives from all others (bottom row) and induces on all others (last column).

**Table 2:** Feedback flux crosstalk

| Ex.\Se. | Ch1 | Ch2 | Ch3 | Ch4 | Ch5 | Ch6 | Ch7 | Sum |
| --- | --- | --- | --- | --- | --- | --- | --- | --- |
| Ch1 | - | 0.095% | 0.014% | 0.017% | 0.004% | 0.003% | 0.001% | 0.133% |
| Ch2 | 0.035% | - | 0.003% | 0.013% | 0.004% | 0.002% | 0.003% | 0.062% |
| Ch3 | 0.049% | 0.002% | - | 0.003% | 0.001% | 0.006% | 0.004% | 0.065% |
| Ch4 | 0.032% | 0.005% | **0.396%** | - | 0.018% | **0.588%** | 0.109% | **1.148%** |
| Ch5 | 0.007% | 0.003% | 0.004% | 0.014% | - | 0.004% | 0.048% | 0.080% |
| Ch6 | 0.013% | 0.008% | 0.054% | 0.041% | 0.008% | - | 0.011% | 0.134% |
| Ch7 | 0.033% | 0.034% | 0.077% | 0.136% | 0.067% | **0.457%** | - | **0.804%** |
| Sum | 0.170% | 0.147% | **0.549%** | **0.224%** | 0.102% | **1.060%** | 0.176% | **2.426%** |

## Acknowledgments

The authors would like to thank Henry Barthelmess for his help modifying the SQUID electronics, Sobhan Sepheri and Edoardo Trabaldo for their help in the lab and Bushra Riaz for fruitful discussions.

This work was financially supported by the Knut and Alice Wallenberg foundation (KAW 2014.0102), the Swedish Research Council (621-2012-3673), the Swedish Childhood Cancer Foundation (MT2014-0007), and Tillväxtverket via the European Regional Development Fund (20201637).

